# A Context-Aware Target Engagement and Pharmacodynamic Biomarker Resource to Accelerate Drug Discovery and Development

**DOI:** 10.64898/2026.04.19.719411

**Authors:** Yuntao Yang, Li Zhao, Seyedmehdi Orouji, Ying Zhu, Rebecca L. Johnson, David S. Maxwell, Ioan Mica, Kaitlyn P. Russell, Bissan Al-lazikani

**Affiliations:** Department of Genomic Medicine, The University of Texas MD Anderson Cancer Center, Houston, TX, USA; Institute for Data Science in Oncology, The University of Texas MD Anderson Cancer Center, Houston, TX, USA; Therapeutics Discovery Division, The University of Texas MD Anderson Cancer Center, Houston, TX, USA; Enterprise Development and Integration, The University of Texas MD Anderson Cancer Center, Houston, TX, USA

## Abstract

Confirming target engagement in tumor experimental models remains a major challenge in oncology drug development. Pharmacodynamic biomarkers can help address this, but few systematic resources link drug targets to candidate biomarkers. We developed TargetTrace, a comprehensive resource to identify and prioritize pharmacodynamic biomarkers across nine key target classes, including transcription factors/cofactors, kinases, phosphatases, ubiquitin ligases, deubiquitinases, acetyltransferases, deacetylases, methyltransferases, and demethylases. Biomarker candidates were gathered from curated molecular interaction resources and refined using external annotations to improve accuracy. For enzyme targets with measurable substrate changes, we applied a two-agent large language model workflow, followed by manual review, to harmonize antibody information from the antibody resources and ensure that the selected biomarkers are measurable with existing laboratory tests. From more than 92,000 input interactions and over 2,300 targets, we compiled 71,323 target-biomarker relationships involving 2,270 potential drug targets, encompassing both transcription factor/cofactor-target gene and enzyme-substrate interactions. Commercial antibodies were available for over 1,400 biomarkers, supporting laboratory validation. This resource provides a structured and reusable resource for systematic identification and prioritization of pharmacodynamic biomarkers in oncology.

## I. Background & Summary

Pharmacodynamic (PD) biomarkers are quantifiable molecular indicators that reflect a drug’s effects on its biological target^1^. Rather than measuring drug exposure alone, PD biomarkers capture the biological consequences of target engagement, thereby linking molecular mechanism to functional response. By assessing biochemical and transcriptional changes following treatment, PD biomarkers provide direct evidence that a therapeutic agent has engaged its intended target and elicited a measurable biological effect. Therefore, they play a central role in modern oncology drug development, supporting mechanism-of-action studies, dose optimization, and early go/no-go decision-making in both preclinical and clinical settings. During earlier drug discovery stages, proximal Target Engagement (TE) biomarkers provide a vital readout to discovery teams to ensure that discovery and optimization remain on track. TE and PD biomarkers thus serve the purpose of robustly maintaining the mechanistic link during drug discovery, development and clinical practice^2^.

Despite their critical importance, systematic resources specifically dedicated to TE/PD biomarkers are still limited. Several existing databases exist for prognostic or patient selection biomarkers. These occasionally include come PD biomarker curation. For example, TheMarker is a curated, publicly available biomarker database contains >16,000 predictive, safety, response or prognostic biomarkers. It contains 218 PD biomarkers spanning 115 drug classes, which were collected through systematic review of scientific literature and U.S. FDA regulatory documents^3^. Similarly, MarkerDB 2.0 is a broad molecular biomarker database constructed through large-scale literature and filtering, and it contains >60,000 biomarkers spanning genetic risks, safety, monitoring and others – but contains only 25 PD biomarkers among its entries^4^. These resources provide limited PD biomarker information and no TE data for mechanistic targets that do not yet have a drug. Moreover, during discovery, monoclonal antibody reagents are typically used by discovery teams to track TE biomarkers. Links to existing antibody reagents is notably absent in the few cases where PD biomarkers are provided in these databases. These limitations are understandable: neither database is dedicated to cover this area and their focus is primarily on response and prognostic biomarkers. Moreover, the coverage is limited to advanced therapeutics and biomarkers and not on supporting discovery stage target engagement studies. As such, they cover a small fraction of potential druggable targets of interest in oncology and beyond.

Given the alarming rate of attrition during drug discovery and development^5^, maintaining a strict audit trail of mechanistic biomarkers is imperative to inform the discovery and development of new treatments. The drug discovery field needs a reliable, large-scale resource of TE/PD biomarkers and available antibody reagents that can be used to detect them in experimental systems. Moreover, it is important that such biomarkers exist within the appropriate biological context being studied. To address these unmet needs, we developed TargetTrace, a large-scale, curated resource of TE/PD biomarkers to support oncology drug development.

**Figure 1a** illustrates the overall workflow. We defined two mechanistically distinct TE/PD biomarker frameworks to cover two major mechanistic classes. The transcriptional framework is designed to identify as proximal a readout as possible for Transcription Factors and Co-factors (TF/COF), here, their downstream target genes were defined as transcriptional PD biomarkers reflecting altered transcriptional activity. The enzymatic framework identifies direct, immediately proximal substrates as enzymatic PD biomarkers reflecting changes in enzyme activity. Enzyme targets were further classified into eight functional groups: kinases, phosphatases, ubiquitin ligases, deubiquitinases, acetyltransferases, deacetylases, methyltransferases, and demethylases.

**Figure 1.**
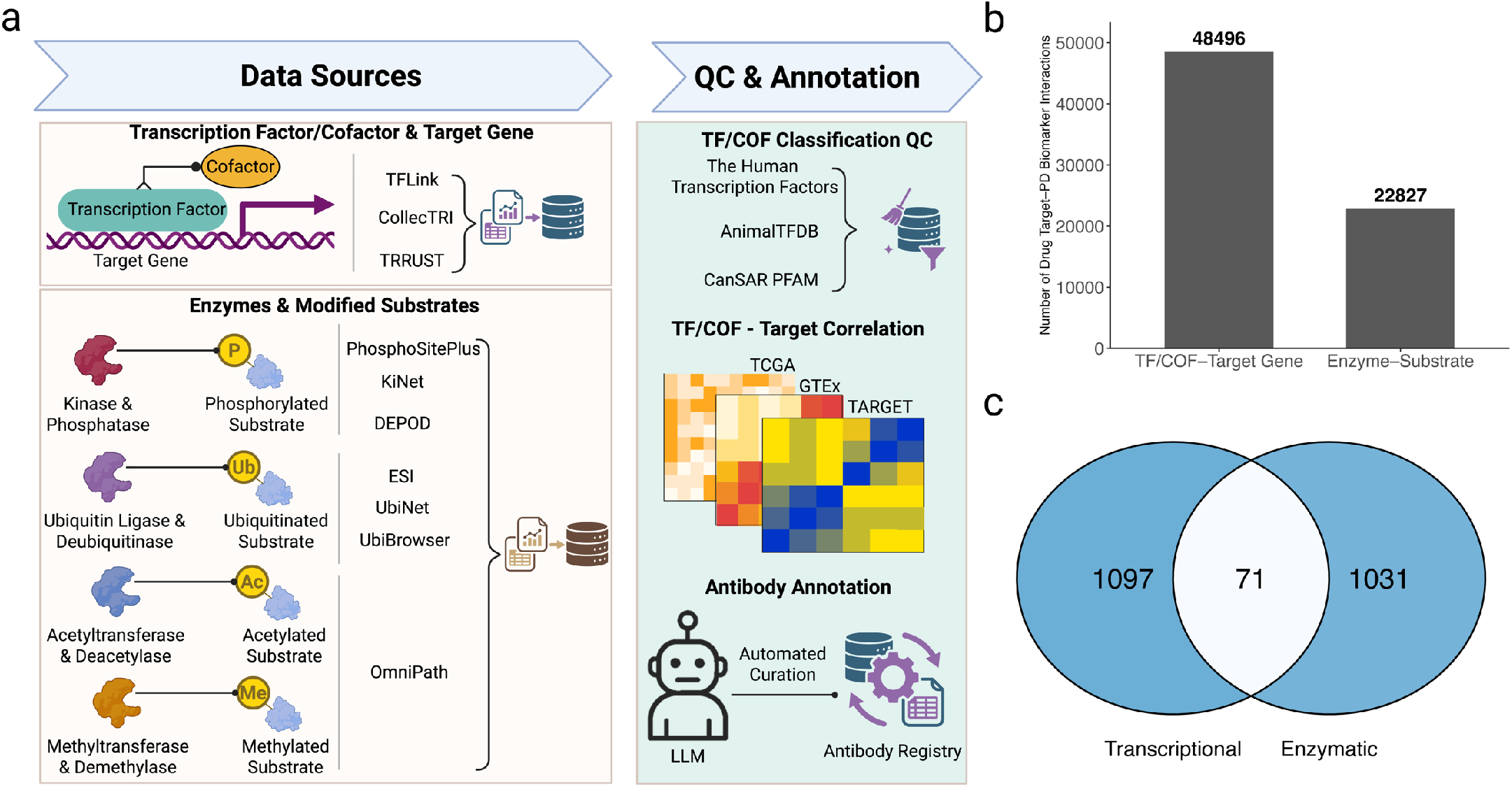
Overview of the TargetTrace construction and landscape. Created with BioRender (https://BioRender.com/rimtre3). **(a)** Schematic of data integration and quality control workflow. **(b)** Total number of curated TF/COF-target gene interactions and enzyme-substrate interactions. **(c)** Number of transcriptional and enzymatic potential drug targets.

During integration of TF/COF-target datasets, we identified proteins incorrectly annotated as TF/COF. To improve accuracy, we cross-referenced interactions with external reference resources and removed misannotated TFs and COFs, thereby enhancing the specificity and reliability of curated transcriptional PD biomarkers. To capture context-specific transcriptional activity, we computed cohort-level correlations between TFs/COFs and their putative target genes, enabling identification of active or repressive relationships in distinct cancer settings. For enzyme targets, we extended beyond substrate curation to improve experimental tractability. A two-agent LLM pipeline was implemented to harmonize antibody metadata from public resources, facilitating identification of validated antibodies for detecting PD biomarkers. Collectively, TargetTrace provides a quality-controlled and experimentally actionable PD biomarker resource, comprising 48,496 TF/COF-target gene interactions and 22,827 enzyme-substrate interactions (**Figure 1b**), spanning more than 2,000 unique transcriptional and enzymatic drug targets for oncology research (**Figure 1c**).

## II. Methods

To construct TargetTrace, we systematically integrated multiple high-quality resources that provide experimentally supported TF/COF-target gene interactions and enzyme-substrate interactions (**Table 1**).

**Table 1.** Sources of drug target-PD biomarker interactions by target class.

| Drug Target Class | Source Database | Evidence / Detection Method | Number of Drug Target-PD Biomarker Interactions |
| --- | --- | --- | --- |
| Transcription Factor/Cofactor | TFLink<sup>6</sup> | Small-scale evidence | 16,634 |
|  | CollecTRI<sup>7</sup> | Public databases; Text mining; Manual curations | 43,857 |
|  | TRRUST<sup>8</sup> | Text mining; Manual curations | 7,874 |
| Kinase | PhosphoSitePlus<sup>9</sup> | Manual curations | 7,404 |
|  | KiNet<sup>10</sup> | Public databases | 8,364 |
| Phosphatase | DEPOD<sup>11</sup> | Manual curations | 934 |
| Ubiquitin Ligase | ESI<sup>12</sup> | Machine learning predictions | 679 |
|  | UbiNet<sup>13</sup> | Public databases; Manual curations | 2,351 |
|  | UbiBrowser<sup>14</sup> | Manual curations; Predictions | 2,891 |
| Deubiquitinase |  |  | 865 |
| Acetyltransferase | OmniPath<sup>15</sup> | Integrate and harmonize multiple resources | 259 |
| Deacetylase |  |  | 18 |
| Methyltransferase |  |  | 68 |
| Demethylase |  |  | 5 |

### 1. TF/COF-Target Gene Interactions

#### Data Acquisition

We integrated TF and COF-target gene interactions from three high-quality resources. First, interactions were obtained from TFLink, contributing 16,634 records from the small-evidence category^6^. Although these interactions do not specify whether the transcriptional effect is activating or repressing, the small-evidence subset represents high-confidence interactions supported by low-throughput experimental methods. In addition, we incorporated 43,857 interactions from CollecTRI, all of which include annotated transcriptional effects and are supported by evidence from public databases, text-mining approaches, and manual curation^7^. Furthermore, 7,874 interactions with defined transcriptional effects were retrieved from TRRUST, supported by text-mining strategies and manual curation^8^.

#### Data Integration & Quality Control

From each data source, we excluded interactions involving self-regulation and non-protein-coding genes. To minimize incorrect annotation of TFs and COFs, interactions were further filtered against three external reference resources: The Human Transcription Factors^16^, AnimalTFDB^17^, and canSAR PFAM (DNA-binding domains)^18^. Only gene products supported by at least one of these resources were retained. Interactions from different databases were harmonized using standardized gene symbols for TFs/COFs and target genes. Identical TF/COF-target gene interactions supported by multiple sources were merged into a single record while preserving all supporting sources as metadata for downstream validation.

When the reported transcriptional effects were inconsistent across two or more sources, the interaction was labeled as “Conflict”; when one source provided a known transcriptional effect and another reported an unknown effect, the known annotation was assigned. Finally, for each target gene, we calculated the number of associated TFs or COFs (*Num_TF_To_Same_Target*) as the count of unique TFs or COFs across all supported interactions, using this metric to quantify redundancy and to prioritize targets with fewer upstream TF/COF inputs as higher-confidence transcriptional PD biomarkers.

### 2. Enzyme-Substrate Interactions

#### Data Acquisition

We incorporated enzyme-substrate interactions from multiple high-quality, curated resources to comprehensively capture major post-translational modification types. Kinase-substrate interactions were obtained from PhosphoSitePlus^9^, contributing 7,404 manually curated interactions supported by experimental studies, and from KiNet^10^, contributing 8,364 interactions derived from public databases. Phosphatase-substrate interactions were retrieved from DEPOD^11^, which provided 936 curated interactions. Ubiquitin ligase-substrate interactions were collected from ESI with 679 machine learning-predicted interactions^12^, UbiNet with 2,351 curated interactions^13^, and UbiBrowser with 2,893 interactions supported by manual curation and predictive evidence^14^; an additional 865 deubiquitinase-substrate interactions were also obtained from UbiBrowser. Acetyltransferase-substrate interactions were retrieved from OmniPath^15^, contributing 259 interactions, along with 18 deacetylase-substrate, 68 methyltransferase-substrate, and 5 demethylase-substrate interactions from the same source. Information on substrate modification sites was included when available, specifically for kinase-substrate interactions from PhosphoSitePlus and KiNet, phosphatase-substrate interactions from DEPOD, and interaction data from OmniPath. In contrast, site annotations were not available for interactions obtained from ESI, UbiNet, and UbiBrowser.

#### Data Integration & Quality Control

Enzyme and substrate identifiers were standardized using official gene symbols to enable cross-database integration. Redundant enzyme-substrate interactions reported by multiple sources were collapsed into a single entry, while all contributing databases were retained as supporting evidence for subsequent analyses. For each enzyme-substrate interaction, we further evaluated exclusivity at both the substrate level and, when available, the modification site level across all curated interactions. *Num_Enzyme_To_Same_Substrate* was defined as the number of unique enzymes associated with a given substrate, and *Num_Enzyme_To_Same_Site* was defined as the number of unique enzymes associated with a specific modification site on that substrate. These metrics were used to prioritize substrates or modification sites with minimal enzymatic redundancy as higher-confidence enzymatic PD biomarkers.

### 3. Cohort-Specific Correlation Analysis for TF/COF-Target Gene Interactions

To identify context-specific transcriptional biomarkers, we performed cohort-specific correlation analyses between each TF or COF and its corresponding target genes using the Pearson correlation coefficient (PCC). Analyses were conducted independently within each cohort derived from TCGA^19^, GTEx^20^, and TARGET^21^. For every TF/COF-target gene interaction in a given cohort, both the PCC value and the associated p-value were calculated. These metrics were used to characterize cohort-specific transcriptional relationships, where positive and negative PCC values suggest putative activating and repressive interactions, respectively.

### 4. Antibody Information Retrieval and Curation using LLM

Antibody information relevant to enzyme-targeted PD biomarkers was obtained from the Antibody Registry, a publicly accessible resource for antibodies^22^. Raw antibody entries were programmatically retrieved in JSON format via the Antibody Registry API. Only antibodies targeting human proteins were retained to ensure translational relevance. Furthermore, to focus on enzyme-mediated modifications, we selected antibodies whose names contained modification-related keywords, including “phospho,” “acetyl,” “methyl,” or “ubiquit,” corresponding to phosphorylation, acetylation, methylation, and ubiquitination, respectively.

Structured extraction of target names, modification types, and modification sites was performed using GPT-5-mini. Prompts were finalized prior to large-scale execution and were not modified during batch processing to ensure reproducibility. A two-agent LLM pipeline was implemented, consisting of an extractor and a validator. In the extraction stage, the extractor was instructed to function as a biomedical text extraction engine and to return exactly one structured JSON object per antibody name. To mitigate formatting inconsistencies and potential hallucination effects inherent to generative models, a separate validator evaluated the extractor’s output using a fixed rule-based prompt introducing new biological inferences or modifying extracted values. The detailed prompt templates for the extractor and the validator are provided in the **Supplementary Methods**. The LLM pipeline was executed in batches of 50 antibody entries per run. Each antibody entry received an overall validity label (“valid” or “invalid”) from the validator. Entries flagged as invalid were manually reviewed and corrected. To further ensure accuracy, a random subset of entries labeled as valid was also independently reviewed and corrected by domain experts.

## III. Data Records

All data records of TargetTrace are deposited through the open access repository on CanSAR^23^.

1. transcriptional_pd_biomarker.txt: This file, provided in TXT format, contains TF/COF-target gene interaction records that define transcriptional PD biomarker interactions. In this framework, the TF/COF represents the potential drug target, and the corresponding target gene represents the transcriptional PD biomarker whose expression change reflects modulation of the target. Each interaction record includes the following features (as indicated by the column headers):
  a. TF_Symbol: Gene symbol of the TF or COF.
  b. TF_EnsemblID: Ensembl ID of the TF or COF.
  c. Is_TF_THTF: Indicates whether the gene is identified as a TF or COF in *The Human Transcription Factors* database.
  d. Is_TF_AnimalTFDB: Indicates whether the gene is identified as a TF or COF in *AnimalTFDB*.
  e. Is_DNA_Binding_CanSAR: Indicates whether the gene contains DNA-binding domains annotated from CanSAR PFAM.
  f. Target_Symbol: Gene symbol of the target gene.
  g. Target_EnsemblID: Ensembl ID of the target gene.
  h. Transcriptional_Effect: Curated transcriptional effect (e.g., activation, repression, conflict, or unknown) describing the direction of transcriptional control.
  i. Num_TF_To_Same_Target: Number of TF or COF regulating the same target gene, used to assess regulatory redundancy or specificity.
  j. Detection_Method: Method used to detect the interaction, as reported by the data source.
  k. Data_Source: Original source of the interaction data.
  l. <CohortID>.COR: Correlation coefficient of the interaction calculated in the specified cohort.
  m. <CohortID>.PVAL: P-value associated with the interaction in the specified cohort.
2. enzymatic_pd_biomarker.txt: This file, provided in TXT format, contains enzyme-substrate interaction records that define enzymatic PD biomarker relationships. In this framework, the enzyme represents the potential drug target, and the corresponding substrate, including site specific modification information when available, represents the enzymatic PD biomarker whose abundance or modification status reflects modulation of enzyme activity. Each interaction record includes the following features (as indicated by the column headers):
  a. Enzyme_Symbol: Gene symbol of the enzyme.
  b. Substrate_Symbol: Gene symbol of the substrate.
  c. Substrate_Modified_Site: Modified site on the substrate protein.
  d. Function_Category: Type of the enzyme.
  e. Data_Source: Original source of the interaction data.
  f. Num_Enzyme_To_Same_Site: Number of enzymes reported to modify the same substrate modification site, used to assess whether the site is uniquely regulated by a given enzyme as a potential PD biomarker.
  g. Num_Enzyme_To_Same_Substrate: Number of enzymes reported to modify the same substrate, used to evaluate whether the substrate is uniquely regulated by a given enzyme as a potential PD biomarker.
3. antibody.txt: This file, provided in TXT format, contains curated antibody information linked to enzymatic PD biomarkers. In this framework, antibodies are used to experimentally measure substrate abundance or site-specific modifications that serve as enzymatic PD biomarkers reflecting modulation of enzyme activity. Each record includes the following features (as indicated by the column headers):
  a. Antibody_Name: Full commercial name of the antibody product.
  b. Target_Name: Gene symbol of the substrate protein recognized by the antibody, corresponding to the enzymatic PD biomarker.
  c. Modification_Type: Type of modification detected by the antibody, including phosphorylation, ubiquitination, acetylation, and methylation.
  d. Modification_Site: Amino acid residue and position recognized by the antibody.
  e. Antibody_Information: Commercial source information for the antibody product, including the vendor name, catalog number, and Research Resource Identifier obtained from the Antibody Registry.

## IV. Technical Validation

All drug target-PD biomarker interactions incorporated into TargetTrace were derived from established, peer-reviewed, and widely adopted community resources to ensure data reliability and biological credibility. To further enhance technical rigor and biological confidence, multiple quality-control procedures were implemented.

### 1. Validation of TF/COF-Target Gene Interactions

More than 900 TF/COF-target gene interactions were removed when the corresponding TF/COF lacked annotation as a transcriptional regulator in any curated reference resource, thereby minimizing potential misclassification. Transcriptional effect annotations distinguishing activation and repression were then systematically compared across supporting data sources. Overall, 88.4% of interactions were consistently annotated as either activation or repression. Interactions with discordant transcriptional effect annotations (n = 679) were retained but explicitly labeled as “Conflict,” rather than excluded, to preserve transparency and allow users to interpret inconsistencies directly. Additionally, 10.2% of interactions were classified as having unknown transcriptional effects (**Figure 2a**). More than 33% of target genes considered transcriptional PD biomarkers were regulated by a single TF/COF, reducing potential confounding from redundant regulatory inputs and enabling clearer assessment of TF/COF target engagement (**Figure 2b**). Following quality control, consistency assessment, and cross-database integration, a total of 48,496 high-confidence TF/COF-target gene interactions were retained. Although CollecTRI contributed the largest proportion of interactions, a subset of TF/COF-target gene relationships overlapped with TFLink and TRRUST, providing independent cross-database support (**Figure 2c**).

**Figure 2.**
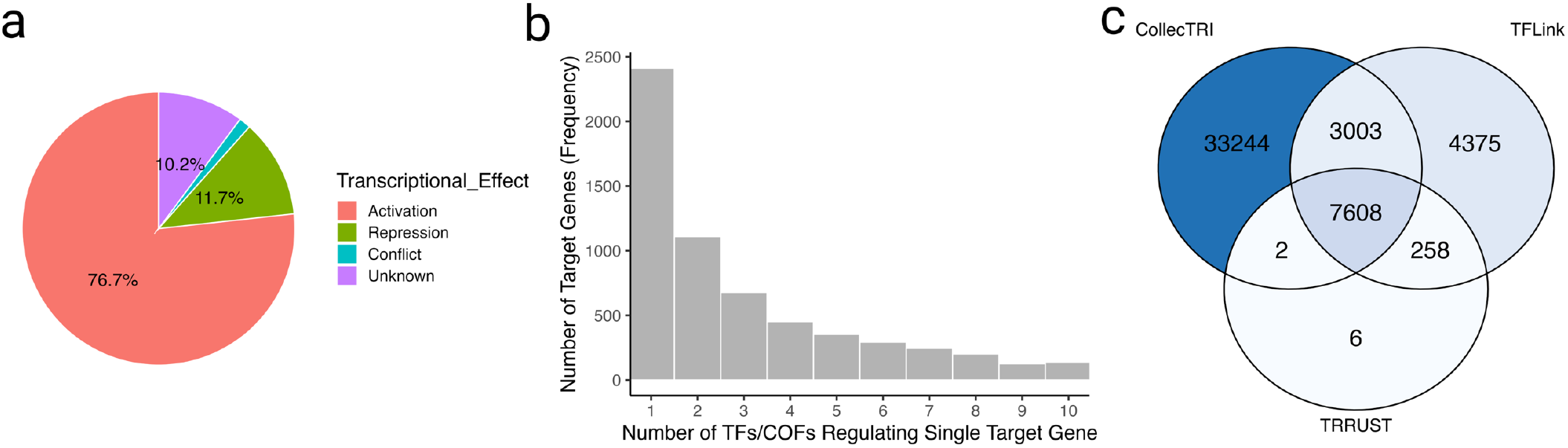
Technical validation of TF/COF-target gene interactions. **(a)** Pie chart summarizing the consistency of transcriptional effect annotations across supporting data sources. **(b)** Histogram showing the distribution of the number of TFs/COFs regulating each target gene (the x-axis is limited to 1 –10 for clarity), highlighting predominant specificity in these interactions. **(c)** Venn diagram depicting the overlap of TF/COF-target gene interactions among source databases.

### 2. Validation of Enzyme-Substrate Interactions

Enzyme-substrate interactions curated in TargetTrace were primarily derived from expert-annotated resources to ensure high-confidence evidence. Most of entries originated from manually curated databases, including PhosphoSitePlus, DEPOD, UbiNet, and UbiBrowser (**Table 1**). To assess the robustness of curated enzyme-substrate interactions, we next examined the degree of overlap across independent data sources. For kinase-substrate interactions supported by multiple independent resources, 83% were shared between KiNet and PhosphoSitePlus, demonstrating strong cross-database concordance. After systematic integration and quality control, a total of 8,615 high-confidence kinase-substrate interactions were retained in the final dataset (**Figure 3a**). Similarly, 79% of ubiquitin ligase-substrate interactions were supported by more than one independent resource, including ESI, UbiBrowser, and UbiNet. Following integration and quality assessment, 2,941 ubiquitin ligase-substrate interactions were retained (**Figure 3b**).

**Figure 3.**
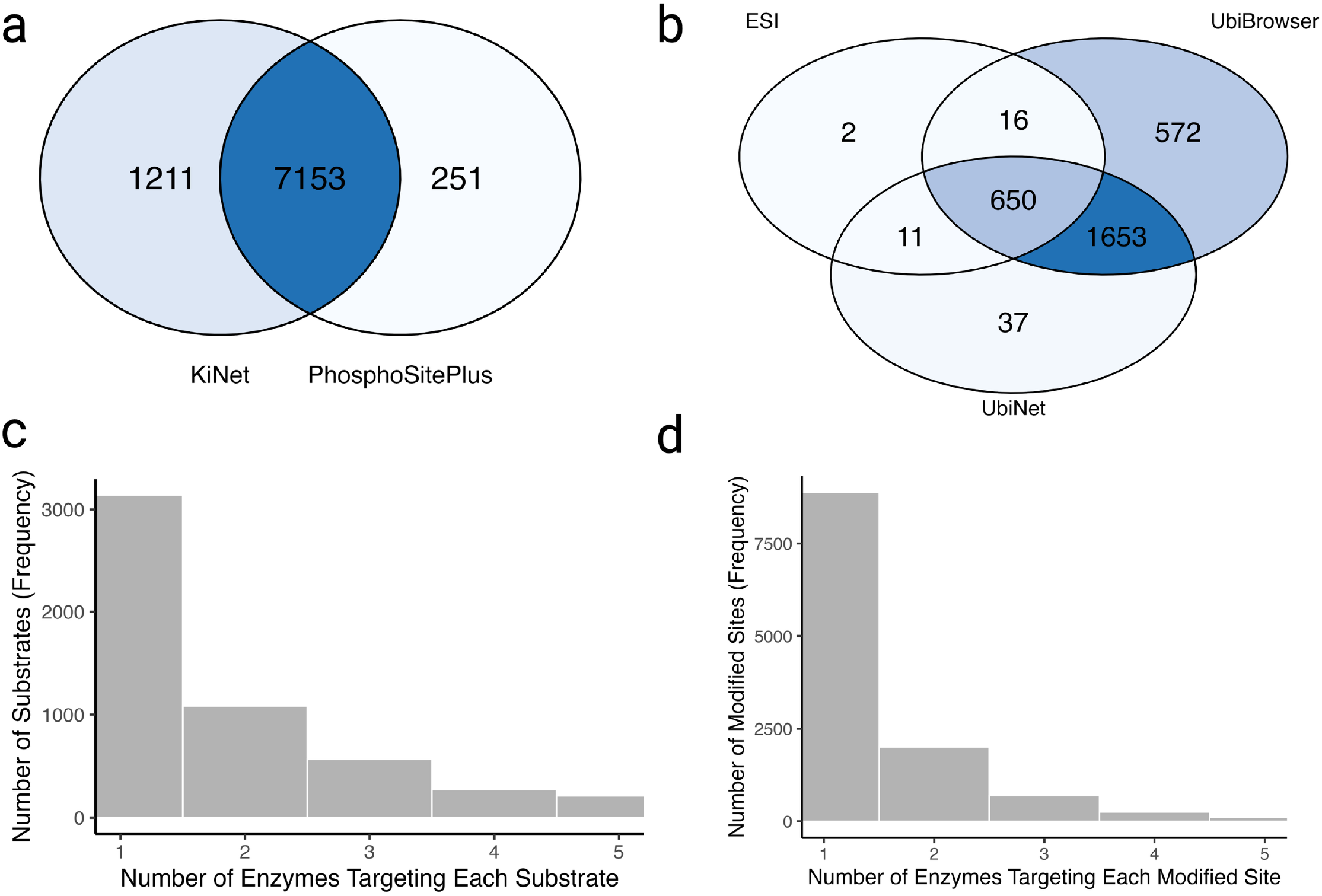
Technical validation of enzyme-substrate interactions. **(a)** Venn diagram showing the overlap of kinase-substrate interactions between KiNet and PhosphoSitePlus. **(b)** Venn diagram illustrating the overlap of ubiquitin ligase-substrate interactions among ESI, UbiBrowser, and UbiNet. **(c)** Histogram showing the distribution of the number of enzymes targeting each substrate, stratified by enzyme class (the x-axis is limited to 1–5 for clarity), to illustrate substrate-level enzymatic specificity and exclusivity. **(d)** Histogram showing the distribution of the number of enzymes targeting each modified site, stratified by enzyme class (the x-axis is limited to 1–5 for clarity), to illustrate site-level specificity and exclusivity.

To further evaluate interaction specificity, we systematically assessed substrates and their modification sites by calculating the number of enzymes targeting each entity within their respective enzyme classes. This stratification revealed that, within specific enzyme types, more than 3,000 substrates were associated with a single enzyme (**Figure 3c**), and over 7,500 modified sites were linked to a single enzyme (**Figure 3d**). These exclusive substrate- or site-level associations represent markers with reduced enzymatic redundancy, thereby improving interpretability of enzyme target engagement and enhancing their potential utility as enzymatic PD biomarkers.

### 3. Validation of Curated Antibody Information

We retrieved 84,141 human antibody entries containing modification-related keywords from the Antibody Registry. These entries were processed through a two-agent LLM pipeline, which only flagged 326 (0.39%) entries as “invalid.” To assess the validity of these flags, we conducted a comprehensive manual review of all 326 entries (**Supplementary Table 1**). Our analysis indicates that the validator successfully identified extractor-induced errors in over 60% of these cases (**Figure 4a**). Specifically, these errors comprised target name inaccuracies (26.1%), modification type discrepancies (20.7%), and modification site errors (13.4%). The remaining flagged entries were attributed to misclassification or scope limitations: 24.9% were false positives, where the validator incorrectly flagged valid entries, and 14.9% were categorized as “out of scope”. These out-of-scope entries primarily involved conditions not addressed by the initial prompt, such as amino acid typos, pan-ubiquitin antibodies, and targets directed at peptide sequences rather than specific amino acid residues. The manual review of invalid entries is available in **Supplementary Table 1**.

**Figure 4.**
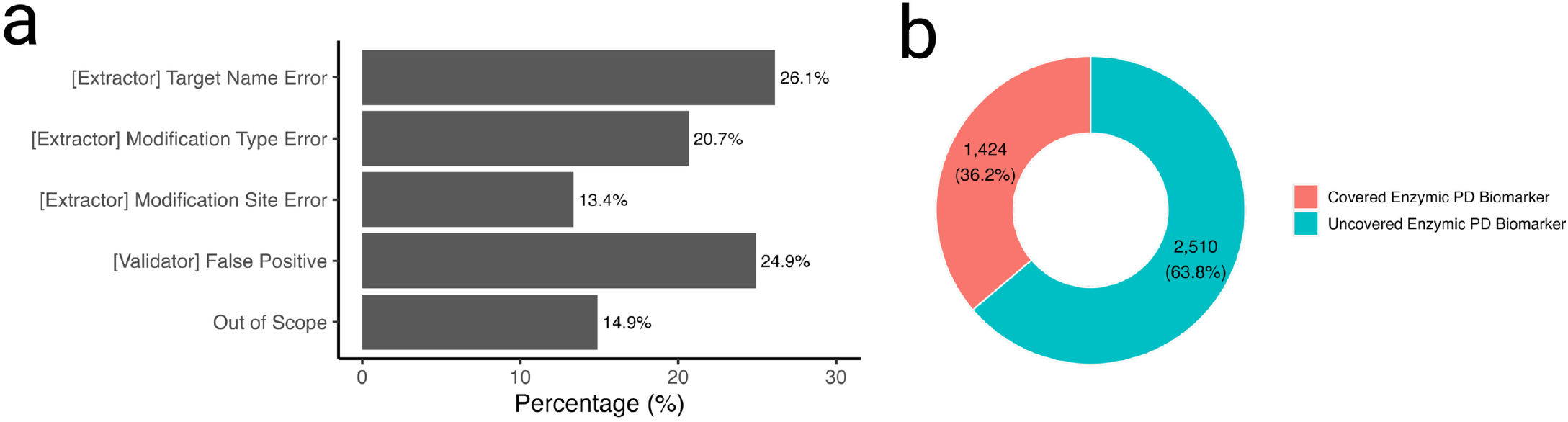
Technical validation of curated antibody information. **(a)** Bar chart showing the percentage of error categories identified within antibody entries flagged as invalid during manual review. **(b)** Donut chart showing the coverage of Enzymic PD Biomarkers by the curated antibody data.

To ensure the high fidelity of our curated dataset, we randomly selected an additional subset of 326 entries from the 99.61% entries that were classified as “valid” by our pipeline and subjected them to independent manual review (**Supplementary Table 2**). This validation confirmed that over 95% of these entries were accurate. The remaining minority of errors (less than 5%) were primarily attributed to the misalignment of modification sites within multi-target interactions and target name errors. While the errors within this sampled subset were manually corrected, it is important to note that the remaining unselected portion of the “valid” dataset, which represents the majority of the repository, is estimated to retain a similar error rate of less than 5%.

Following the manual review and correction of both the “invalid” and “valid” subsets, the final curated dataset resulted in 59,594 high-quality antibody entries. This final set provides coverage for 1,424 (36.2%) of the identified enzymic PD biomarkers (**Figure 4b**). Ultimately, the high accuracy rate within the “valid” subset and the successful identification of errors among the limited “invalid” flags, coupled with targeted manual corrections, resulted in a high-fidelity repository that effectively maps enzymatic PD biomarkers to reliable antibody reagents.

## V. Usage Notes

To utilize the PD biomarker data in TargetTrace, users should begin by searching for the drug target of interest, classified as either a TF/COF or an enzyme, to identify corresponding PD biomarkers. For drug targets classified as TF/COFs associated with multiple transcriptional PD biomarkers, selection should be prioritized by favoring candidates reported across multiple data sources and those with the lowest *Num_TF_To_Same_Target* values, which indicate exclusive transcriptional relations. Users should then evaluate *Transcriptional_Effect* annotations to classify interactions as activation or repression and assess correlations specific to the study cohort. For enzyme targets associated with multiple enzymatic PD biomarkers, prioritization should favor candidates reported across multiple data sources and those with the lowest *Num_Enzyme_To_Same_Site* or *Num_Enzyme_To_Same_Substrate* values, indicating exclusive enzymatic relations. Users should consult antibody annotations to identify existing antibodies for these enzymatic PD biomarkers. Users are encouraged to carefully assess antibody quality, specificity, and suitability for their intended application. In scenarios where multiple exclusive enzymatic PD biomarkers are available, testing them simultaneously is recommended to ensure mechanistic robustness and internal validation of target engagement.

## VI. Data Availability

The data comprising TargetTrace are freely accessible via the CanSAR open-access repository^23^.

## VII. Code Availability

No custom code or algorithms were used to generate or process the data described in TargetTrace.

